# Core components of endosomal mRNA transport are evolutionarily conserved in fungi

**DOI:** 10.1101/467787

**Authors:** Jessica Müller, Thomas Pohlmann, Michael Feldbrügge

## Abstract

Active movement of mRNAs by sophisticated transport machineries determines precise spatiotemporal expression of encoded proteins. A prominent example discovered in fungi is microtubule-dependent transport via endosomes. This mode of transport was thought to be only operational in the basidiomycete *Ustilago maydis*. Here, we report that distinct core components are evolutionarily conserved in fungal species of distantly related phyla like Mucoromycota. Interestingly, orthologues of the key RNA-binding protein Rrm4 from the higher basidiomycete *Coprinopsis cinerea* and the mucoromycete *Rhizophagus irregularis* shuttle on endosomes in hyphae of *U. maydis*. Thus, endosomal mRNA transport appears to be more wide-spread than initially anticipated.

**Highlights:**

‐ Core transport components Upa1 and Rrm4 are conserved in different fungal phyla
‐ Components of the Rrm4 machinery were most likely secondarily lost in ascomycetes
‐ Upa1 from *Microbotryum lychnidis-dioicae* is functional in *U. maydis*
‐ Rrm4 orthologues from Basidio- and Mucoromycota shuttle in hyphae of *U. maydis*

## Introduction

mRNA trafficking is important in a variety of cellular processes (Eliscovich and Singer, 2017). A widespread mechanism is the active transport of mRNAs along actin or microtubules (Mofatteh and Bullock, 2017). Key factors are RNA-binding proteins (RBPs) that recognize cargo mRNAs and interact with accessory RBPs to form larger ribonucleoprotein complexes (mRNPs). These are linked to molecular motors (Niessing et al., 2018).

A well-studied example is actin-dependent transport of *ASH1* mRNA towards the distal pole of daughter cells during cytokinesis in *Saccharomyces cerevisiae*. The key RBPs She2p and She3p bind cargo mRNAs cooperatively and She3p interacts with the myosin motor Myo4p during transport (Edelmann et al., 2017; Niessing et al., 2018). Orthologues of She2p are only found in Saccharomycetaceae and absent in the closely related *Candida albicans*. The latter contains a She3p orthologue that is needed for hyphal growth (Elson et al., 2009) suggesting conservation of actin-dependent mRNA transport in this group of ascomycetes.

Endosomal mRNA transport along microtubules is well-studied in *U. maydis* and essential for efficient unipolar growth of infectious hyphae (Haag et al., 2015; Vollmeister et al., 2012). The endosomal machinery mediates bulk distribution of mRNAs as well as transport of specific mRNAs like all four septin mRNAs. Co-translational transport of septin mRNAs supports assembly of heteromeric complexes on endosomes. Septin heteromers are transported towards the growth pole to form higher-order filaments (Baumann et al., 2014; Zander et al., 2016).

The key RNA-binding protein Rrm4 binds with its three N-terminal RNA recognition motifs (RRMs; Fig. 1A) thousands of target mRNAs in their 3’ UTR (Olgeiser et al., 2018). Rrm4 contains two C-terminal MademoiseLLE domains (MLLE domains) for interaction with PAM2-like motifs. Two of these are present in Upa1 that links Rrm4-containing mRNPs via its FYVE domain to endosomes (Fig. 1B; Pohlmann et al., 2015). Transported mRNPs are stabilized by the scaffold protein Upa2 that contains four PAM2 motifs for interaction with the poly(A)-binding protein Pab1 (Fig. 1A-B, Jankowski et al., 2018). In addition, Upa2 contains a functionally important GWW motif for interaction with currently unknown transport factors. In essence, Rrm4, Upa1 and Upa2 constitute functional core components of endosomal mRNP transport (Fig. 1B).

**Fig. 1:**
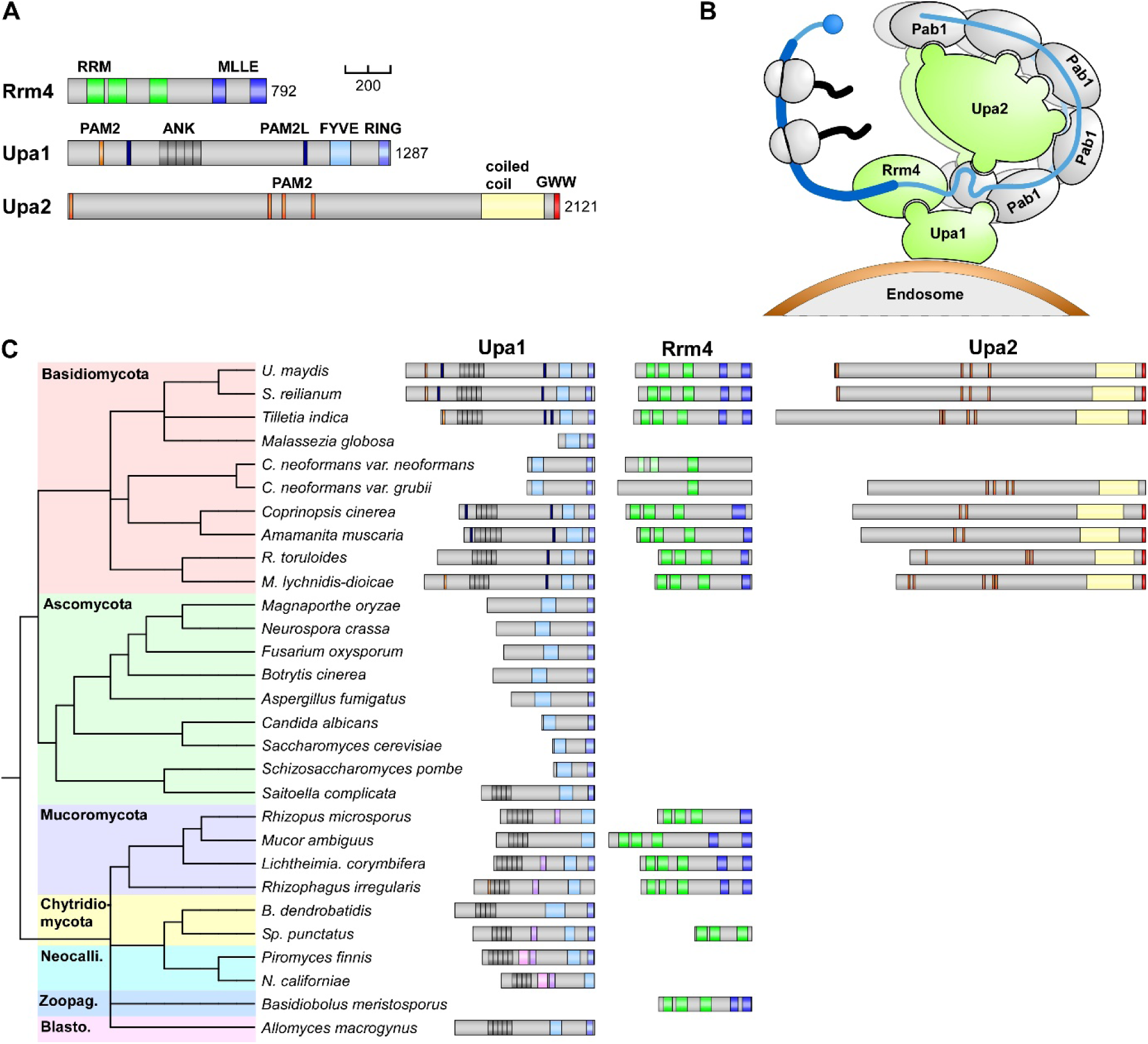
Components of endosomal transport are conserved in distinct fungal phyla. (A) Schematic representation of core components of endosomal mRNA transport: the MLLE domain-containing protein Rrm4 as well as the PAM2 motif-containing proteins Upa1 and Upa2 drawn to scale (bar, 200 amino acids; green, RRM domain; blue, MLLE domain; orange, PAM2 motif; dark blue, PAM2-like motif; dark grey, Ankyrin repeats; light blue, FYVE domain; blue, RING domain; yellow, coiled coil region; red, GWW). Some orthologues of Upa1 contain a DysFN and DysFC domain indicated in rose and lilac. (**B**) Schematic representation of the core components (green) of endosomal mRNA transport. (**C**) Schematic representation of orthologues of Upa1, Rrm4 and Upa2 in representative organisms of different fungal phyla (Neocalli., Neocallimastigomycota; Zoopag, Zoopagomycota; Blasto, Blastocladiomycota. The phylogenetic tree was generated by applying the phyloT tool using the NCBI taxonomy (no evolutionary distances). If no protein is depicted, a clear orthologue could not be identified. Accession numbers are listed in Supplementary Table S5. S., *Sporisorium*; C., *Cryptococcus*; R., *Rhodotorula*; M., *Microbotryum*; B., *Basidiobolus*; Sp., *Spizellomyces;* N., *Neocallimastix*.

## Results and Discussion

### Core components of the endosomal mRNA transport machinery are conserved in fungi

To address whether endosomal mRNA transport along microtubules is conserved in fungi, we performed a phylogenetic analysis of core components Rrm4, Upa1 and Upa2. All three components are present in Basidiomycota, with orthologues in Ustilaginomycotina, Puccionomycotina and Agraricomycotina (Fig. 1C). Interestingly, Upa2 is only absent in few Basidiomycota that also lack clear Rrm4 orthologues like *Malassezia globosa* and *Cryptococcus neoformans* suggesting secondary loss. An exception is *C. neoformans var. grubii* containing a weakly conserved Upa2-like protein. However, it lacks the functionally important GWW motif. Consistently, the potential Rrm4 orthologue lacks MLLE domains for interaction with endosomal Upa1 (Fig. 1C).

Importantly, Rrm4 and Upa1 orthologues are also found in Mucoromycota indicating that the key RBP of endosomal transport and its adaptor are not restricted to basidiomycetes. Potential orthologues of Rrm4 are found in Chytridiomycota and Zoopagomycota, but are most likely absent in Ascomycota, Neocallimastigomycota and Blastocladiomycota (Fig. 1C). The endosomal linker protein UmUpa1 contains a C-terminal FYVE domain for binding endosomal lipids and a PAM2-like motif for interaction with Rrm4 (Pohlmann et al., 2015). Notably, these features are conserved in those Basidiomycetes containing Rrm4 and Upa2 orthologues. In most ascomycetes, orthologues are shorter missing the N-terminal Ankyrin repeats for protein interaction. An interesting exception is *Saitoella complicata*, which combines characteristics of Ascomycota and Basidiomycota (Nishida et al., 2011). Furthermore, N-terminally extended Upa1 orthologues are found also in all other fungal clades suggesting that the Ankyrin repeat-containing version is primordial. In essence, the core machinery consisting of Rrm4, Upa1 and Upa2 appears to be conserved in Basidiomycota. The presence of Upa1 and Rrm4 in distant phyla like Mucoromycota suggests that fundamental principles of endosomal transport might be wide-spread in fungi.

### Distantly related orthologues of Rrm4 shuttle in *Ustilago maydis*

To test the functionality of Rrm4 and Upa1 orthologues, we expressed Gfp-tagged versions in *U. maydis* (see Methods). Since we had to codon-optimise distantly related genes, we focused on few examples. Initially, we asked whether the orthologues of Rrm4 and Upa1 were able to complement characteristic growth defects of disturbed mRNA transport, i.e. increased number of bipolar growing hyphae. The closely related orthologues from *Sporisorium reilianum* were able to complement (Fig. 2A-D) and exhibited a similar shuttling like their *U. maydis* counterparts (Fig. 2E-F).

**Fig. 2.**
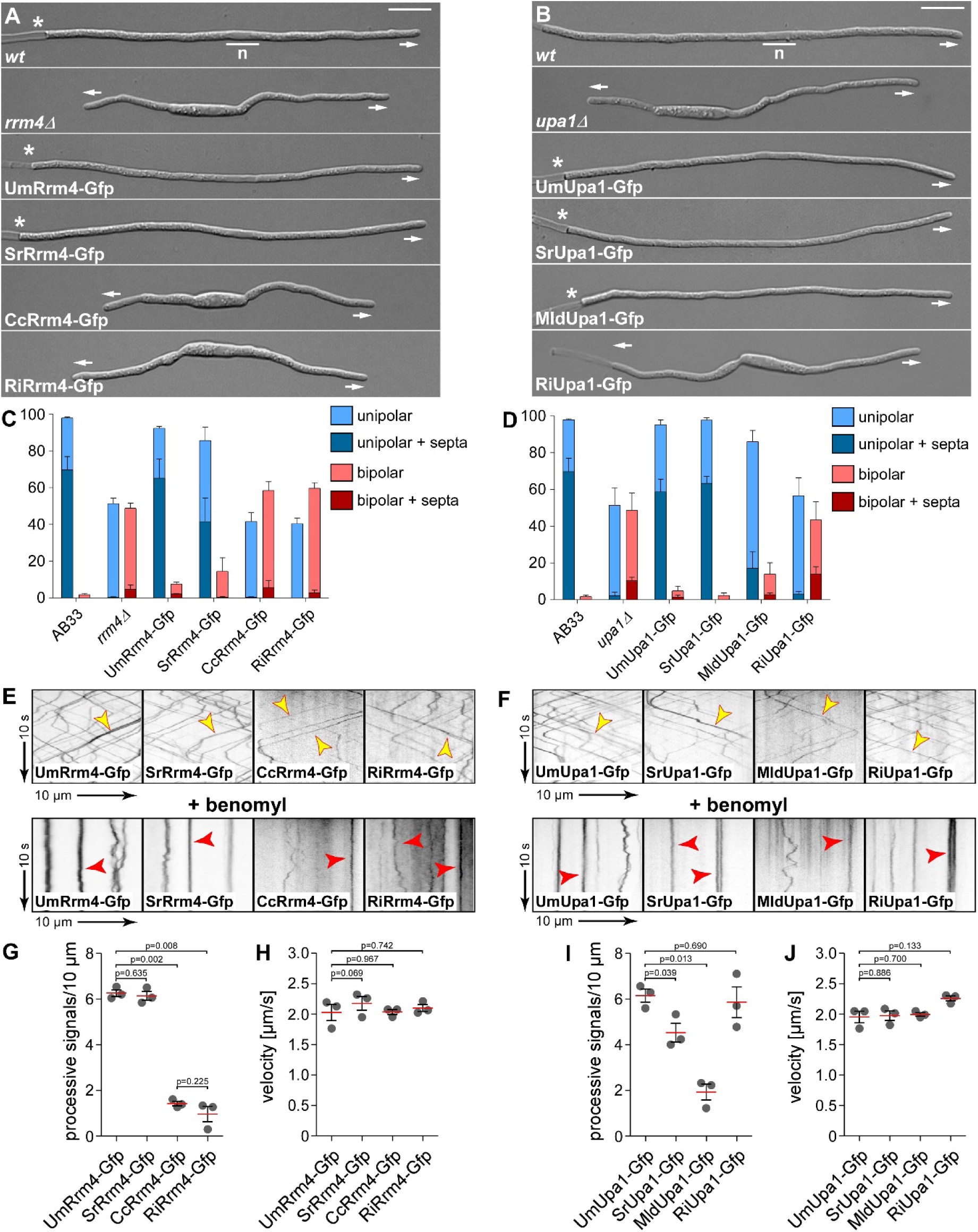
Orthologues of Rrm4 and Upa1 shuttle microtubule-dependently on endosomes. (**A, B**) Growth of AB33 hyphae (6 h.p.i.; growth direction is marked by arrows; n, nucleus; asterisks, empty sections). Um, *Ustilago maydis*; Sr, *Sporisorium reilianum*; Cc, *Coprinopsis cinerea*; Ri, *Rhizophagus irregularis*; Mld, *Microbotryum lychnidis-dioicae*. (**C, D**) Quantification of hyphal growth (6 h.p.i): unipolarity, bipolarity and septum formation were quantified (error bars, s.e.m.; n = 3 independent experiments, for each experiment >100 hyphae were analysed per strain; note that septum formation is given relative to the values of unipolar or bipolar hyphae set to 100%). (**E, F**) Kymographs of AB33 hyphae expressing different orthologues of Rrm4 or Upa1 C-terminally fused to eGfp (6 h.p.i.); genetic background as indicated (arrow length on the left and bottom indicate time and distance, respectively). Processive movement is visible as diagonal lines (top lane, yellow arrowheads), static signals after disruption of microtubule cytoskeleton upon treatment with benomyl (bottom lane, red arrowheads). (**G, I**) Processive signals per 10 pm of hyphal length (data points representing mean from n = 3 independent experiments, with mean of means, red line, and s.e.m.; paired two-tailed Student’s t-test, for each experiment at least 10 hyphae were analyzed per strain). (**H, J**) Velocity of fluorescent signals (velocity of tracks with > 5 μm processive movement; data points representing means from n = 3 independent experiments, with mean of means, red line, and s.e.m.; paired two-tailed Student’s t-test; for each experiment at least 30 signals per hypha were analyzed out of 10 hyphae per strain).

Studying the subcellular localisation of more distantly related Upa1 orthologues revealed that all versions moved bidirectionally throughout hyphae with velocities resembling microtubule-dependent movement of transport endosomes (Video 1; Fig. 2F, J; Jankowski et al., 2018). Upa1 from *Microbotryum lychnidis-dioicae* did shuttle less frequently in comparison to the others, but it retained functionality (Fig. 2C, 2I).

Analysing the subcellular localisation of Rrm4 orthologues showed that Rrm4 from *S. reilianum* shuttled comparably to Rrm4 from *U. maydis* (Fig. 2E, G, H). Importantly, also Rrm4 orthologues from *Coprinopsis cinerea* and *Rizophagus irregularis* shuttled, although the number of processive units was reduced (Video 1). All Rrm4 fusion proteins exhibited characteristic velocities of endosomal transport (Fig. 2H, J) and processive movement could be inhibited by the microtubule-destabilizing drug benomyl (Fig. 2E, F, bottom). Therefore, we conclude that these distant orthologues hitchhike on transport endosomes arguing for a conserved mechanism.

## Conclusions

Core components of the endosomal mRNA transport machinery appear to be conserved in various fungal phyla. This suggests that important features of endosomal transport might be more wide-spread than currently anticipated. Although no clear orthologues of Rrm4, Upa1 and Upa2 are found in plants and animals, recent findings also suggest a link between endosomes and mRNA trafficking in these organisms (Konopacki et al., 2016; Yang et al., 2018, Béthune et al., 2018).

## Methods

### Plasmids, strains and growth conditions

The coding regions of SrUpa1 and SrRrm4 were amplified by PCR from *S. reilianum* gDNA. The coding regions of MldUpa1, CcRrm4, RiUpa1 and RiRrm4 were optimized according to context-dependent codon usage (Zarnack et al., 2006) and ordered from IDT (Skokie, IL, USA). All coding regions were C-terminally fused to *eGfp* (enhanced Gfp, Clontech; SrRrm4 was fused to *eGfp* and Tap tag) and integrated in the native loci of *rrm4* and *upa1* in AB33rrm4Δ and AB33upa1Δ, respectively. All integration events were verified by Southern analysis. All genes are under control of the native *U. maydis* promoters. Strains and plasmids are listed in Supplemental Tables S1-S4. Detailed growth conditions and induction of hyphal growth have been described previously (Baumann et al., 2012; Brachmann et al., 2001; Baumann et al., 2015).

### Microscopy, image processing and image analysis

The microscopy setup has been described previously (Jankowski et al., 2018; Pohlmann et al., 2015). Fluorescence micrographs are displayed inverted. For quantification of hyphal growth, cells were imaged and scored for unipolarity, bipolarity and septum formation 6 h.p.i. (hours post induction). Three independent experiments were conducted with more than 100 cells per strain. For analysis of signal number and velocity of Gfp fusions, videos were recorded with 150 frames and an exposure time of 150 ms. Kymographs were generated and analyzed using Metamorph (Version 7.7.0.0, Molecular Devices, Seattle, IL, USA). Processive signals were counted manually. Three independent experiments (n = 3) were conducted with at least 10 hyphae per strain. The mean velocity was determined for processive signals (movement >5 μm) using Metamorph. Three independent experiments (n = 3) were conducted with at least 10 hyphae per strain. Benomyl treatment has been described elsewhere (Jankowski et al., 2018).

### Phylogenetic analysis and bioinformatics

Bioinformatic analysis was conducted as described previously (Jankowski et al., 2018). Accession numbers of the proteins used for phylogenetic analysis and complementation experiments are listed in Supplemental Table S5.

## Acknowledgements

We acknowledge L. Olgeiser for generating SrRrm4-GfpTT as well as Dr R. Kellner and lab members for discussion and reading of the manuscript. We are grateful to U. Gengenbacher for excellent technical assistance. The work was funded by grants from the Deutsche Forschungsgemeinschaft to MF (FE448/10-1 DFG-FOR2333; DFG-CRC1208 and CEPLAS EXC1028).

---

**Legend to Video 1:**
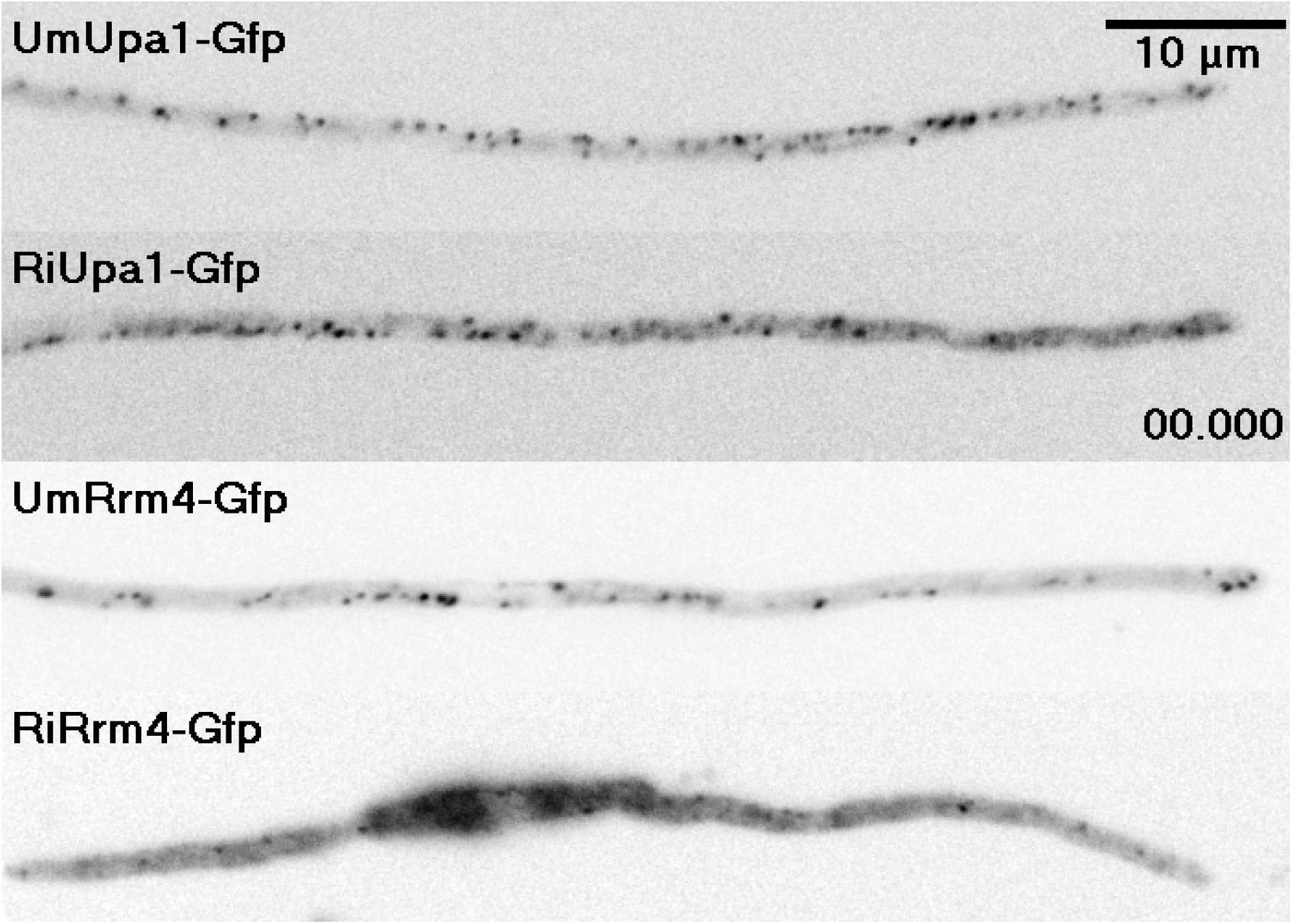
Orthologues of Rrm4 from higher basidiomycetes and mucoromycetes shuttle on endosomes in hyphae of *U. maydis*. Composite video showing UmUpa1-Gfp in comparison to RiUpa1-Gfp (top two lanes) and UmRrm4-Gfp and RiRrm4-Gfp (bottom two lanes). All proteins are shuttling within hyphae, with RiRrm4-Gfp showing a prominent localization in the cytoplasm (scale bar, 10 μm; timescale in seconds, 150 ms exposure time, 101 frames, 6 frames/s display rate; MP4 format, 1427 kb).

